# High-throughput single-cell sorting by stimulated Raman-activated cell ejection

**DOI:** 10.1101/2023.10.16.562526

**Authors:** Jing Zhang, Haonan Lin, Jiabao Xu, Meng Zhang, Xiaowei Ge, Chi Zhang, Wei E. Huang, Ji-Xin Cheng

## Abstract

Single-cell sorting is essential to explore cellular heterogeneity in biology and medicine. Recently developed Raman-activated cell sorting (RACS) circumvents the limitations of fluorescence-activated cell sorting, such as the cytotoxicity of labels. However, the sorting throughputs of all forms of RACS are limited by the intrinsically small cross-section of spontaneous Raman scattering. Here, we report a stimulated Raman-activated cell ejection (S-RACE) platform that enables high-throughput single-cell sorting based on high-resolution multi-channel stimulated Raman chemical imaging, *in situ* image decomposition, and laser-induced cell ejection. The performance of this platform was illustrated by sorting a mixture of 1 μm polymer beads, where 95% yield, 98% purity, and 14 events per second throughput were achieved. Notably, our platform allows live cell ejection, allowing for the growth of single colonies of bacteria and fungi after sorting. To further illustrate the chemical selectivity, lipid-rich *Rhodotorula glutinis* cells were successfully sorted from a mixture with *Saccharomyces cerevisiae*, confirmed by downstream quantitative PCR. Furthermore, by integrating a closed-loop feedback control circuit into the system, we realized real-time single-cell imaging and sorting, and applied this method to precisely eject regions of interest from a rat brain tissue section. The reported S-RACE platform opens exciting opportunities for a wide range of single-cell applications in biology and medicine.

**Significance statement:** Image-guided single-cell sorting is a potent tool in diverse biological applications. Current microfluidic cell sorting methods encounter challenges in handling smaller cells and are not applicable to tissue sections. To address these challenges, we have developed a stimulated Raman-activated cell ejection (S-RACE) platform, which is the first demonstration of single-cell ejection coupled with coherent Raman scattering. S-RACE allows label-free chemical imaging guided cell sorting through multispectral stimulated Raman scattering (SRS) imaging, on-the-fly image analysis, and laser-induced cell ejection. Versatile applications of S-RACE to a wide range of samples, such as polymer particles, single-live bacteria, single-live fungus, and tissue sections, are demonstrated.

## Introduction

Cell sorting is indispensable for characterizing a heterogeneous cell population from various perspectives, such as chemical, structural, and genomic analyses (1–3). Current cell sorting techniques include flow cytometry, laser microdissection, cell picking, and microfluidics (4, 5). Among this array of techniques, fluorescent and magnetic labeling are commonly used for sorting target cells, whereas the exogenous labels may potentially induce cytotoxicity and disrupt cellular functions. Additional challenges such as the lack of specific labels and susceptibility to photobleaching limit the utility of labeling-based cell sorting. In contrast, label-free cell sorting methods rely on cellular properties like morphology, deformability, and chemical content. However, the morphological and mechanical attributes may not exhibit a direct correlation with biological states, thus reducing the sorting specificity (6). For example, quantitative phase imaging could map the volumetric distribution of the refractive index but is difficult to identify specific cells (7, 8).

Raman spectroscopy possesses the capacity to surpass the constraints faced by the above regimes. By detecting inelastic photon scattering (9), Raman spectroscopy could characterize the endogenous chemical content of single cells and is capable of probing cell metabolic activity (10). By integrating Raman spectroscopy with cell sorting methodologies, including flow (11, 12), optical tweezer (13, 14), dielectrophoresis (15, 16), and cell ejection (17, 18), a plurality of biomedical applications were achieved (19). Raman-activated cell sorting (RACS) has found extensive use in microbiology for isolating functional individuals from a community. It has been illustrated to differentiate antibiotic-resistant (20–22) or functional microbes that possess specific pathways in a complex environment such as the human gut (21) or natural ecosystems (23–25). Other cell types like mammalian and fungal cells can also be characterized and sorted using RACS (16, 26, 27). Among the RACS techniques, Raman-activated cell ejection (RACE) has been proven especially powerful in high-precision sorting of smaller cells. RACE is based on laser-induced forward transfer, a method widely used in material transfer (28). In RACE, the specimens are first placed on coverslips with a laser-absorbing coating material. A pulsed laser then acts on the target location to ablate the coating, providing forward momentum to eject the cells to the collector for downstream analysis (19), for example, linking single-cell phenotype and genotype of microorganisms sampled from the natural environment (24). Despite its versatility, current RACS methods have a low throughput due to the small cross-section of spontaneous Raman scattering. Typical integration time for a single Raman measurement ranges from 15 and 60 seconds per spectrum for biological samples to obtain a sufficient signal-to-noise ratio (29).

Here, we demonstrate stimulated Raman-activated ejection (S-RACE) to achieve automated high-throughput cell sorting. In coherent Raman microscopy, 10^4^∼10^5^ signal enhancement can be achieved compared to spontaneous Raman scattering (30, 31). Coherent Raman microscopy employs two pulsed laser beams to probe coherent vibrations in a sample. Both stimulated Raman scattering (SRS) and coherent anti-Stokes Raman scattering (CARS) have been combined with microfluidics for high-throughput cell imaging (32–36). Recently, two coherent Raman-activated cell sorting studies were reported, one in the C-H stretching region (37) and the other in the fingerprint region (400-1800 cm^-1^) by an FT-CARS spectrometer (38). Despite the high throughput, these microfluidic-based methods encounter challenges in handling smaller cells (<3 μm) (37), and unstable flow caused by bubbles and/or debris in the microfluidic channel (39). Apart from cell detection and sorting, SRS microdissection and sequencing was recently reported for *in situ* laser microdissection of tissue slices and downstream DNA and RNA sequencing (40).

Our S-RACE platform integrates multicolor SRS imaging, online image processing, and a lab-built ejection module to enable high-throughput image-based cell sorting. This is the first demonstration of single-cell ejection guided by coherent Raman microscopy. We achieved a high yield of approximately 95% and a high purity level of around 98% for a mixture of 1.0-μm polymer beads, with a throughput of approximately 14 events per second (eps). Additionally, we demonstrated fast identification and sorting of lipid-rich *R. glutinis* cells from a mixture with *S. cerevisiae*, and confirmed the result by quantitative PCR amplification of the ITS2 region. A notable feature of our platform is its live cell sorting capability, a pivotal component for isolating and purifying cells with specific functions while ensuring their viability. Successful cell recovery was observed for both bacteria (*E. coli*) and fungus (*S. cerevisiae*). Furthermore, by harnessing a comparator circuit for communication between imaging and laser ejection, we achieved real-time SRS-guided sorting of single polymer beads, live cells, and regions of interest in a brain tissue slice. The S-RACE platform promises various biomedical applications, including metabolic engineering (41), precise diagnosis (29), and bacterial therapies (42).

## Results

### S-RACE platform

Our S-RACE system includes a multispectral SRS microscope and a laser ejection module (**Figure 1A**). The SRS microscope was described in our previous work (41). Detailed setup can be found in **Figure S1**. A multicolor SRS stack was collected by scanning the interpulse delay between the spectrally chirped pump and Stokes beams. The ejection module (**Figure 1B**) consists of a 532-nm 1.0-ns pulsed laser, an ejection coverslip, and a collector assembled in a sandwich-like manner. The ejection coverslip is coated with a thin layer of titanium dioxide (TiO_2_) as the dynamic release layer (43). For the polymer microparticle mixture sorting, the sample is first loaded onto the coverslip with TiO_2_ coating, followed by an air-drying step in preparation for ejection. During ejection, the TiO_2_ coating absorbs energy from the 532-nm laser and generates gas pressure pushing the targeted microparticle away. The “ejected” microparticle is then collected by the bottom collector, which is fabricated by coating a layer of polydimethylsiloxane (PDMS) on a standard coverslip. For live cell sorting, an additional thin layer of agarose gel is introduced on the TiO_2_ coating, mitigating potential mechanical damage during the ejection process (18). In addition, a complementary layer of agarose is introduced at the bottom of the collector to safeguard the cells from any impact during the landing process (18, 44).

**Figure 1.**
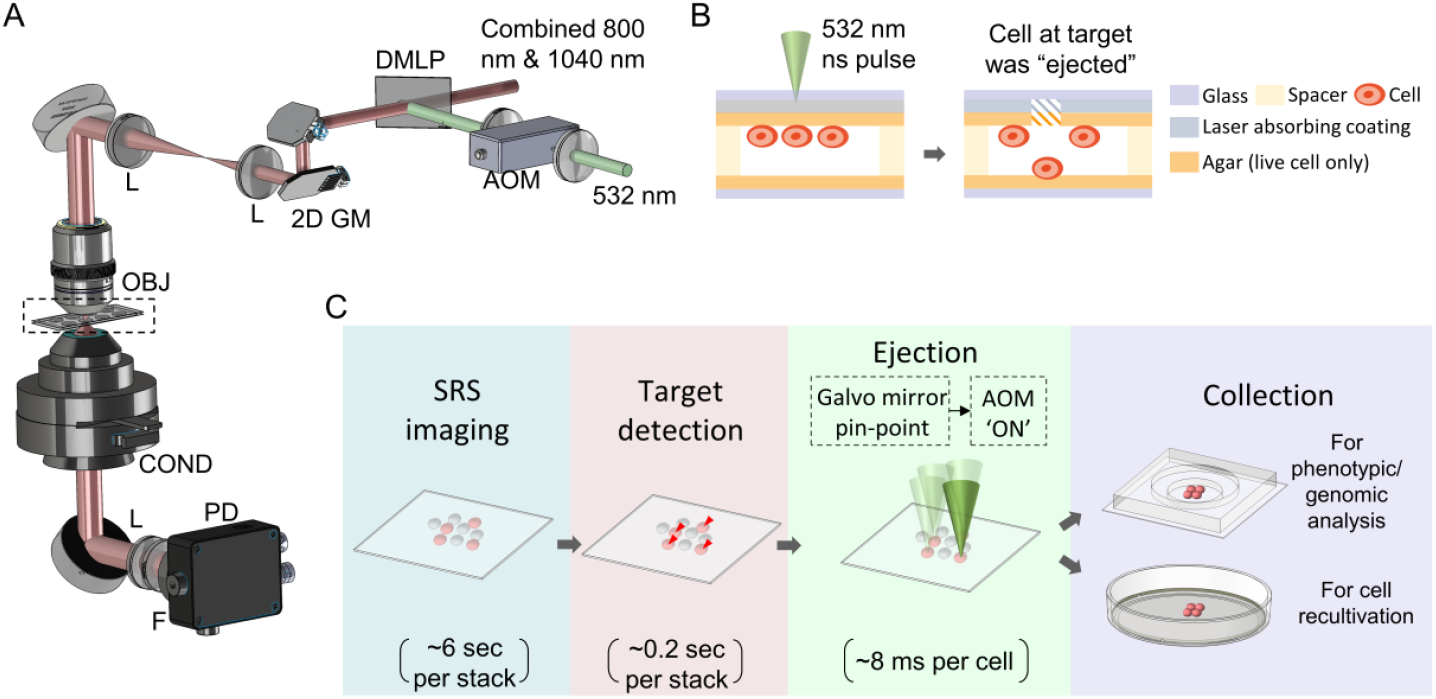
Optical diagram and workflow of S-RACE. **A**. Optical diagram of S-RACE. AOM: acousto-optic modulator. DMLP: long pass dichroic mirror. 2D GM: 2D galvo mirrors. L: lens. OBJ: objective. COND: condenser. F: filter. PD: Photodiode. **B**. Sketch of the ejection process. The left diagram is a zoom-in of the sample in the dashed box in A. **C**. Workflow of S-RACE. Each image stack contains 300x300x4 pixels.

Metals such as gold, metal oxides, and polymer materials have been used as the dynamic release layer in laser-induced forward transfer (LIFT) systems for cell ejection or tissue dissection (43). Among various coatings tested in this study, TiO_2_ showed the best performance in both SRS imaging and laser ejection (**Figure S2**). The TiO_2_-coated coverslip (∼150 µm thick) is fabricated by magnetron sputtering to 4-nm thickness for 2-minute sputtering time. **Figure S3A** shows that TiO_2_ coating has minimal background in SRS imaging and no interference with the SRS signal of polymer beads. In contrast to TiO_2_ coating, commonly used Au-coated coverslips contributed a significant thermal background to the SRS image (**Figure S3B**). The UV-Vis spectrum of the TiO_2_-coated coverslip shows an absorbance peak at 532 nm (**Figure S3D**), which facilitates the laser ejection. To characterize the spatial resolution of ejection, bead clusters dried on the TiO_2_-coated coverslip were used as a test bed (**Figure S3E**). For the 60X objective, the ejection spot diameter is 4.20 ± 0.74 µm. For the 40X objective and underfilled objective back aperture, the ejection spot size is 6.19 ± 1.00 µm. The ejection spot is larger than the optical resolution, probably due to the domino effect of ejection. The test was performed with a minimum laser energy guaranteeing a successful ejection without photodamage. Specifically, the laser power used was lower than 1.5 mW before the objective, and the energy of the laser pulse was less than 1 µJ. We further studied the impact of the axial focal position on the ejection efficiency (**Figure. S3F**). It was found that maximal efficacy was reached when the objective was focused on the TiO2 coating, which is about 1 micron above the beads.

The workflow of S-RACE is shown in **Figure 1C**. For each field of view (FOV), SRS images are first collected at 10 μs pixel dwell time. An SRS stack of 300 by 300 pixels and 4 wavenumbers takes ∼6 seconds. Subsequently, microparticles/cells within the FOV are identified by spectral analysis. For the discrimination of target objects in the context of 4-color SRS image involving two types of polymer microbeads, the target detection step consumes ∼0.2 seconds. For each targeted microparticle/cell, 2D galvo mirrors were employed to precisely position the green laser on the object. An acousto-optic modulator (AOM) was then activated to emit the 532-nm, 4-kHz laser onto the coating. The targeted object was then pushed away from the ejection coverslip and received by the collector coverslip. With AOM as a fast pulse picker, single-pulse ejection is achieved and each ejection takes ∼8 ms, which is sufficient to stabilize 2D galvo mirrors pinpoint and compensate for laser repetition rate. Multiple FOVs were stitched to a larger FOV by moving the sample stage. After sorting all the targeted cells, downstream phenotypic and/or genomic analysis, e.g., sequencing and proteomics, could be applied to the cells in the collector.

For live cell ejection and cultivation, an agarose dish is used as a collector. After incubating the collector with sorted live cells at an appropriate temperature (30°C for fungus and 37°C for bacteria), single colonies can be observed and recovered after 24 to 48 hours, as illustrated in the following sections.

### S-RACE performance evaluated with mixtures of polymer microbeads

We tested S-RACE performance using a polymer bead mixture (polystyrene (PS), with red fluorescence; poly (methyl methacrylate) (PMMA), without fluorescence; both 1.0 μm in diameter). Four wavenumbers (2860, 2905, 2950, and 2994 cm_-1_) representing the Raman signatures of PS (2905 cm^-1^) and PMMA (2950 cm^-1^) were selected (**Figure S4A**). This polymer beads test was performed with a laser excitation from the top configuration (**Figure S3E**), which features a small focus size. A composite image of two-color SRS before ejection is shown in **Figure 2A**. All the beads were classified into two types (PS or PMMA) based on the workflow in **Figure S4B**, and PS beads were targeted for sorting. The composite SRS image with classified beads color-labelled is shown in **Figure 2B**. To further enhance sorting purity, PS beads (the targeted class) that have neighboring PMMA beads located closer than 2.5 µm (identified as “clustered” PS) were excluded from sorting. The post-ejection identity of the beads is visualized in **Figure 2C**. In the zoom-in image of **Figure 2B**, two white arrows highlight PMMA beads situated adjacent to a PS bead. Both PMMA beads were retained following the ejection of all the targeted PS beads. Averaged spectra of beads before and after ejection were shown in **Figure 2D**, confirming the beads classification. During experiments, we observed bright spots remaining on the coverslip after ejection (**Figure 2C**). The spectra of these bright spots (**Figure S4C**) were distinguishable from those of PS or PMMA (**Figure S4A**), indicating that these bright spots might have been caused by the deformations in the TiO_2_ coating (45, 46).

**Figure 2.**
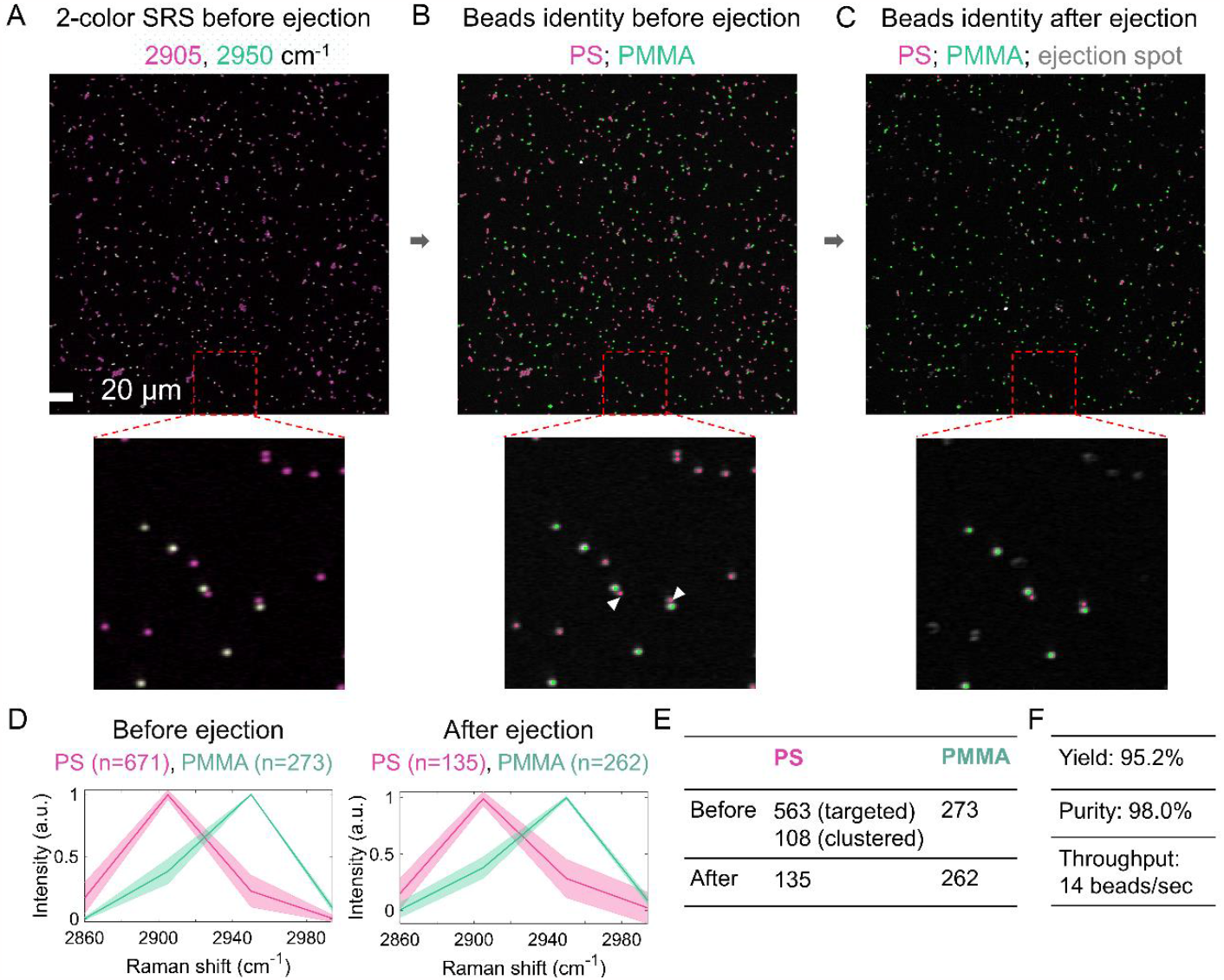
Evaluation of S-RACE performance by polymer microbead mixtures (polystyrene (PS), with red fluorescence; poly(methyl methacrylate) (PMMA), without fluorescence; both 1.0 μm in diameter). **A**. 4-color SRS image before ejection. **B**. Beads identified based on A. White arrows in zoom-in image mark the detected 2 PS beads which excluded from ejection. **C**. Beads identify after ejection. **D**. Multicolor SRS of polymer beads before and after ejection. Shaded error bar: standard deviation. **E-F**. Quantification of ejection performance.

The quantification of ejection performance is presented in **Figure 2E-F**. Our S-RACE achieved 95% yield, 98% purity, and ∼14 events per second (eps). Given that the PS beads possessed red fluorescence, we were able to quantify the PS beads in the collector using wide-field fluorescence (**Figure S4D**). This analysis revealed the presence of 292 PS beads in the collector, with a collection rate of ∼55%. Experiments conducted using different bead exclusion criteria demonstrated that stricter exclusion criteria resulted in higher purity (**Figure S5**).

### S-RACE is applicable to live cells

To validate the biocompatibility of our S-RACE platform in sorting single live cells, we conducted tests on *S. cerevisiae, C. albicans*, and *E. coli*, representing both eukaryotic and prokaryotic cells. Hydrogels, polymers, and aqueous media layers have been reported in printing viable cells (18, 47). In this study, to protect live cells from heat and mechanical damage process, we adapted the protocols from Liang (18) and Hong (48) to prepare an agarose layer on TiO_2_ coating. The thickness of the agarose gel is 6.25 ± 0.95m based on estimation under a microscope. As shown in **Figure 3A-B**, individual *S. cerevisiae* cells on the agarose gel were identified based on bright-field imaging. The image of the agarose layer after ejection is shown in **Figure 3C**. The agarose plate with the sorted cells was then sent to a 30 °C incubator, and ∼27 colonies were observed after ∼44 hours (**Figure 3D**). The recovery rate of single *S. cerevisiae* cell ejection was ∼31%. The control group, in which non-cell areas were ejected, did not lead to any colony growth on the receiving agar plate. Similar tests were performed on *C. albicans* (**Figure 3E-H**) and GFP-labeled *E. coli* (**Figure 3I-L**). Cultivation recovery rates were ∼23% and ∼21% for *C. albicans*, and *E. coli*, respectively. Recultivated GFP-labeled *E. coli* colonies were observed (**Figure 3L**). These results established the coating condition for live cell ejection.

**Figure 3.**
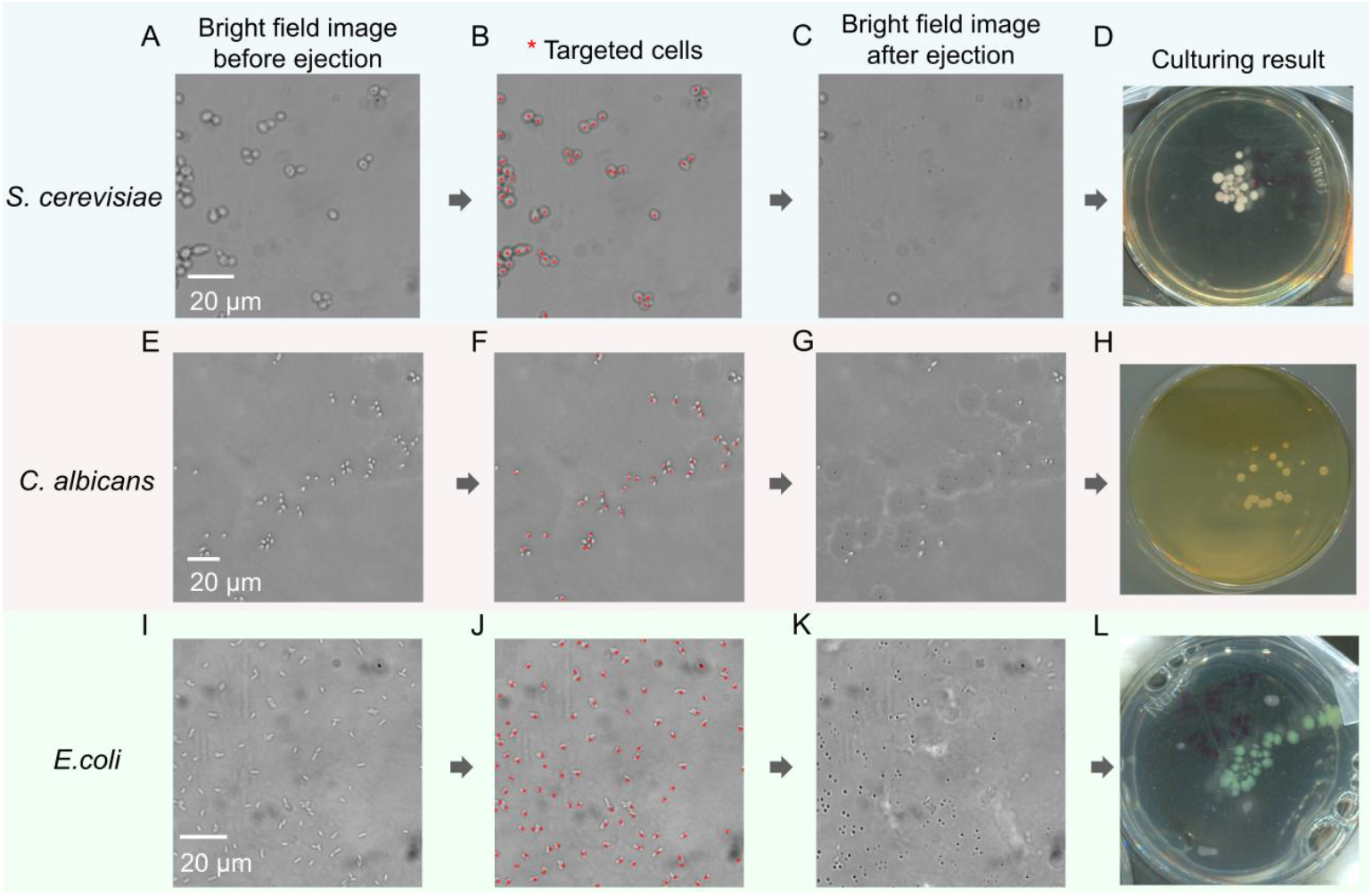
Live cell ejection (*S. cerevisiae, C. albicans*, and *E*.*coli*). **A-D**. Bright field image guided sorting of *S. cerevisiae* (cultivation recovery rate ∼31%). **E-H**. Bright field image guided sorting of *C. albicans* (cultivation recovery rate ∼23%). **I-L**. Bright field image guided sorting of GFP-labelled *E. coli* (cultivation recovery rate ∼21%).

To assess the impact of SRS laser radiation on cell viability during the S-RACE of live cells, time-lapse imaging was performed to visualize *E. coli* growth after SRS laser radiation (**Figure S6**). Three different SRS laser radiation levels were chosen: no radiation, low radiation (pump 24 mW, Stokes 50 mW), and medium radiation (pump 24 mW, Stokes 100 mW). The medium radiation was more stringent compared to the experimental condition. *E. coli* cells were dropped onto a 1% agarose pad after sampling from liquid culture, and sandwiched with a top coverslip. The cells were kept in an enclosed incubator stabilized at 30°C. For all three laser radiation levels, cell growth and colony formation were observed. The growth rates, calculated by fitting the growth curve of the cell colony areas, were not significantly different across the three radiation levels. This result confirms the biocompatibility of the SRS laser radiation.

We then performed S-RACE of two types of live cells: *S. cerevisiae* (**Figure 4A-F**) and GFP-labeled *E. coli* (**Figure 4G-L**). **Figure 4A** shows the schematic of the ejection module. **Figure 4B** shows SRS spectra in the C-H stretching region of the cells. Individual *S. cerevisiae* cells were identified based on single-frame SRS images at 2935 cm_-1_ (signal-to-background ratio ∼4, **Figure 4C-D**). SRS image after ejection is shown in **Figure 4E**. Bright spots in **Figure 4E** were probably caused by the deformed agarose layer and/or deformed TiO_2_ coating. The spectra of bright spots after ejection are shown in **Figure S7**, different from the spectra of cells or the image background. Recultivated cells were transferred to culture tubes after ∼40 hours of cultivation on an agarose plate. The culture medium with ejected cells became turbid, indicating successful cell growth, whereas the medium of the control group remained clear. For GFP-labeled *E. coli*, the collector used was composed of a thin agar layer (∼60 µm) and a standard coverslip. This design enables the visualization of sorted cells in the collector. **Figure 4I-K** shows the image before and after ejection. After cell ejection, a wide-field fluorescence image of the sorted GFP-labeled *E. coli* in the collector confirmed that most of the cells remained in good shape (**Figure 4L**).

**Figure 4.**
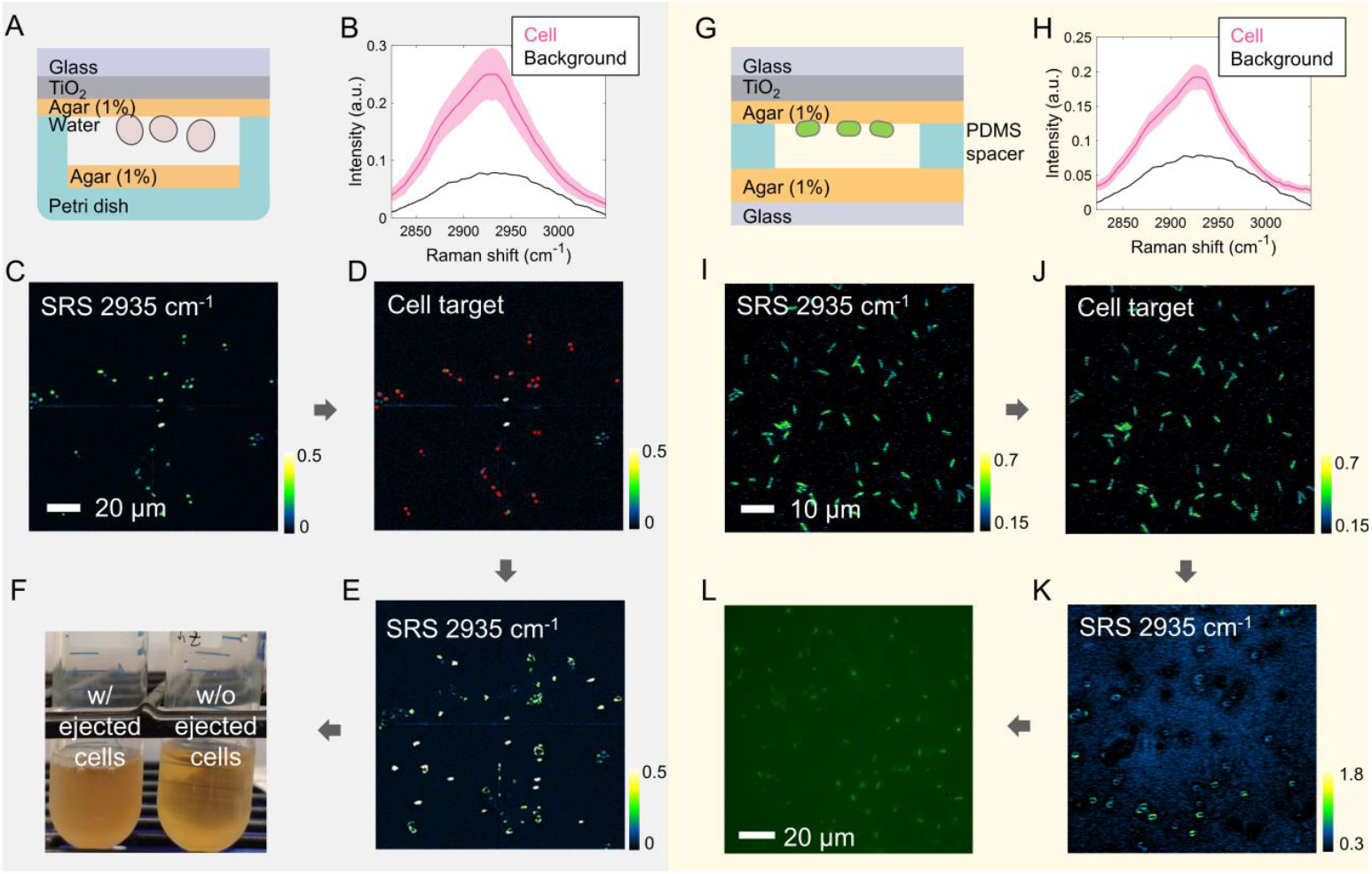
S-RACE and recovery of live cells (*S. cerevisiae* and GFP labelled *E. coli*). **A**. Schematic of the ejection module used for *S. cerevisiae* ejection. **B**. SRS spectra of *S. cerevisiae* and background. **C**. single-color SRS image of *S. cerevisiae* before ejection. **D**. Cell identified based on C. **E**. Single-color SRS image of the same FOV as D after ejection. **F**. Picture of culture tube after 48 hours. **G**. Schematic of the ejection module used for *E. coli* ejection. **H**. SRS spectra of *E. coli* and background. Shaded error bar: standard deviation. **I**. Single-color SRS image of *E. coli* before ejection. **J**. Cell identified based on I. **K**. Single-color SRS image of the same FOV as J after ejection. **L**. Wide-field fluorescent image of collected *E. coli* on the bottom agar.

### S-RACE of lipid-rich *R. glutinis* from cell mixture followed by qPCR identification

To show the utility of our S-RACE platform to sort target cells from a mixture, we tested lipid-rich *Rhodotorula glutinis* (*R. glutinis*) cells mixed with *S. cerevisiae. R. glutinis* has been identified as oleaginous yeast and can contain up to 70% lipids in its dry-weight biomass (49, 50). In contrast, *S. cerevisiae* only has ∼6% lipids in its biomass (49). Lipid-rich yeasts including *R. glutinis* have been valuable models for sustainable biofuel production. Using multicolor SRS, the lipid content in individual yeast cells can be quantified. Lipid-rich intracellular aggregates found in the SRS image had a spectrum similar to that of glycerol trioleate (**Figure 5A-B**). While the cell body part has a similar spectrum as peptone, a standard protein sample (**Figure 5B**), the lipid channel after background subtraction in **Figure 5C** showed the SRS signal at 2851 cm^-1^ is contributed by lipid (51).

**Figure 5.**
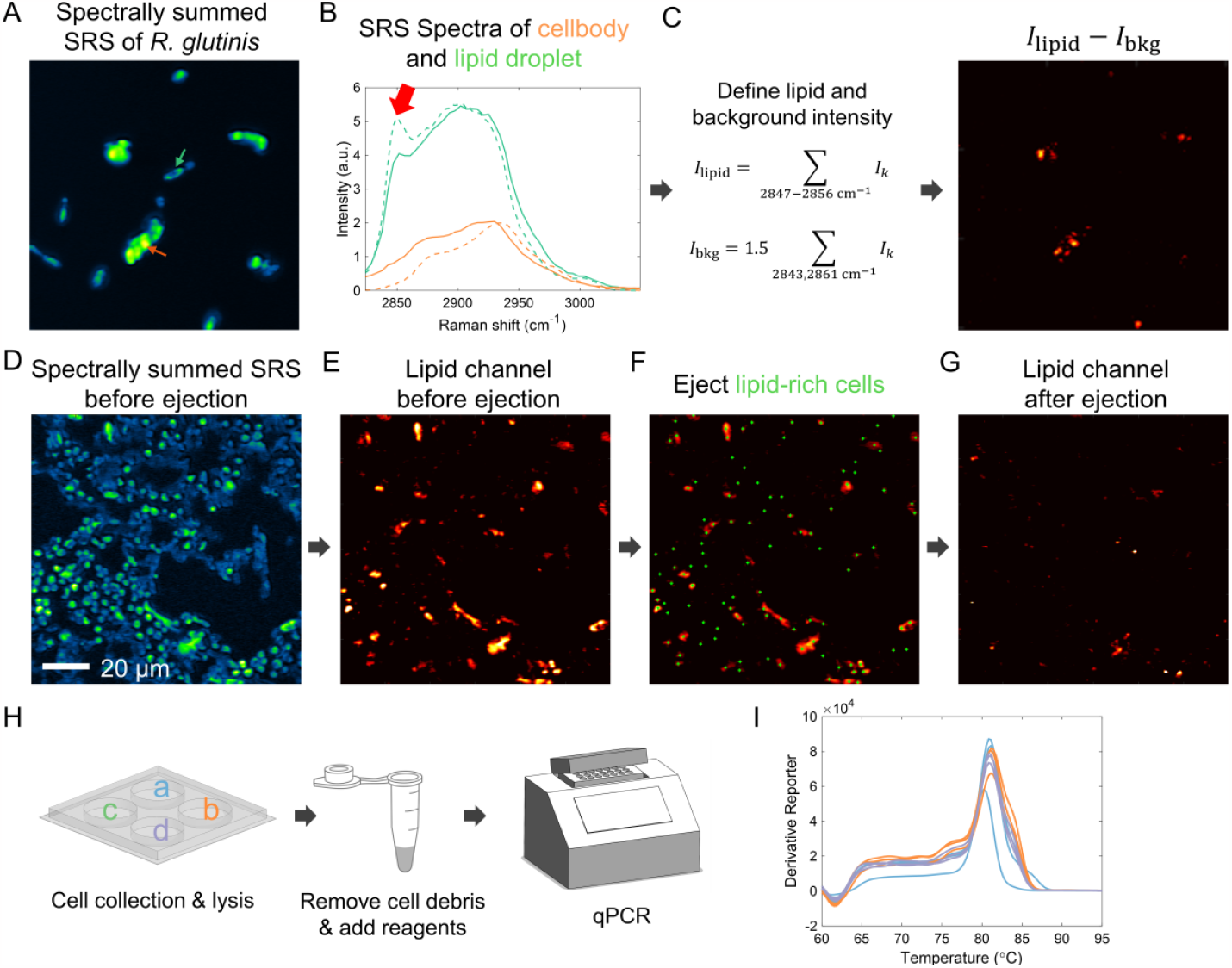
S-RACE of lipid-rich *R. glutinis* from mixture with *S. cerevisiae* and qPCR identification. **A**. Spectrally summed hyperspectral SRS of *R. glutinis*. **B**. SRS Spectra of cell body (yellow solid line) and lipid droplet (green solid line), with two standard samples (glycerol trioleate: green dashed line; peptone: yellow dashed line). The red arrow denotes the signature peak of lipid. **C**. Left: definition of lipid and background intensity. Right: lipid channel of the same FOV in A. Lipid channel presents SRS signal at around 2850 cm^-1^ with linear background subtracted. **D**. Spectrally summed 5-color SRS image of *R. glutinis* and *S. cerevisiae* mixture before ejection. **E**. Lipid channel of the same FOV in D. **F**. Location of lipid-rich cells found by automatic program based on lipid channel in E. **G**. Lipid channel after ejection. **H**. Sample preparation steps for qPCR amplification. Left: schematic of ejected cell collector (a-d represent 4 independent trials); middle: qPCR vial; right: qPCR machine. **I**. Melt curve of the PCR amplicons (different color represents different trials, and each trial produced 3 replicates).

**Figure 5D** shows the spectral summation of a 5-color SRS image of a mixture of *R. glutinis* and *S. cerevisiae* (ratio ∼1:1). The lipid-rich cells were subsequently identified from the lipid channel (**Figure 5E-F**). The cells were sorted with the bottom configuration in Figure S3E. After sorting the lipid-rich cells, the lipid channel has a much reduced intensity (**Figure 5G**), which confirms the successful ejection of lipid-rich cells. The sorted cells were collected by a customized collector made with coverslip and PDMS (**Figure 5H**). Four independent trials were performed (presented as a-d in **Figure 5H**). Each well received sorted cells from 8-12 SRS FOVs. The throughput is ∼11.9 cells per second.

To confirm the identity of the sorted cells are *R. glutinis* we targeted, quantitative polymerase chain reaction (qPCR) was performed on the collected cells. The second internal transcribed spacer (ITS2) in the nuclear ribosomal DNA was used as the target sequence. The workflow of qPCR amplification preparation is shown in **Figure 5H**: after DNA extraction (52), the supernatant containing resuspended DNA was used for qPCR, using primers from a previous study (53). Each well produced 3 replicates in qPCR amplification, and each replicate contained ∼16% of the total DNA content in this well. The amplifications of the collected cell contents present a peak of around 81.3°C in the melt curve (**Figure 5I**), consistent with the amplification results of *R. glutinis* pure culture without ambiguous peaks (**Figure S8B**). We would also like to note that the cell number per well estimated from qPCR amplification is lower than the number of ejection due to the cell loss during the transfer process and the presence of larger cells that necessitate multiple ejections (**Figure S8 C-D**). In conclusion, these results demonstrate that S-RACE successfully sorted specific cell populations based on their phenotype/functions.

### S-RACE with opto-control allows for real-time sorting of cells in culture and regions of interest in a tissue slice

Seamlessly integrating a real-time precision opto-control (RPOC) system with our S-RACE platform yielded a real-time cell sorting approach. The RPOC utilizes a closed-loop feedback control circuit for laser manipulation with a fast response of sub-microsecond (54). This innovation enables imaging-identifying-sorting to occur within a single pixel during laser scanning, bringing opportunities for higher precision and efficiency. The concept of real-time imaging sorting is illustrated in **Figure 6A**. The front panel of the comparator circuit is shown in **Figure 6B**. During laser scanning, the SRS signal carrying chemical information from the sample was sent to the comparator circuit. For an SRS signal higher than the preset threshold, the comparator circuit will command AOM to rapidly couple the 532 nm laser, which subsequently ejects the targeted object residing in the current pixel. Limited by the repetition rate of 532 nm laser (16.6 kHz), 70 µs dwell time was applied. To show the utility of this real-time imaging-sorting regime, we sorted 1-µm polymer beads, single cells, and tissue sections based on their SRS images. For polymer beads, the SRS image (2950 cm^-1^) before sorting is shown in **Figure 6C** (top). Active pixels were set by thresholding the SRS image, and each polymer bead contained 4-6 active pixels for best sorting performance. Most of the polymer beads were successfully sorted based on their SRS intensity. For live *S. cerevisiae*, SRS images before and after real-time imaging sorting are shown in **Figure 6D**. After ∼48 hours, six *S. cerevisiae* colonies were observed in the petri dish with sorted cells and no colony growth was observed in the control group.

**Figure 6.**
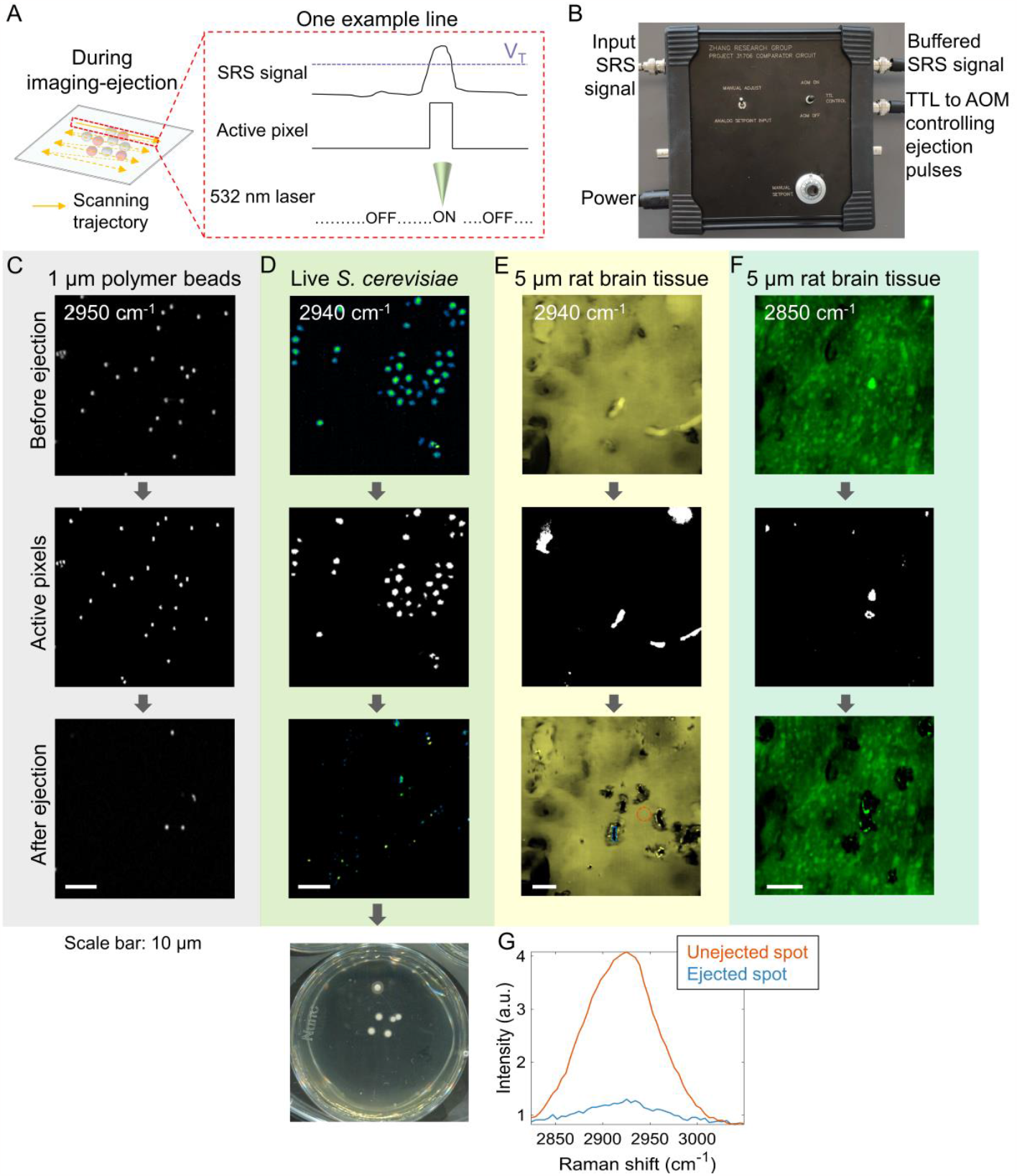
Real-time imaging and sorting of cells in culture and in tissue environment with a real-time precision opto-control (RPOC) system. **A**. An illustration of real-time imaging-sorting with RPOC technology. **B**. Front panel of the comparator circuit box with ports used in this study. **C**. Real-time imaging-sorting of 1 µm polymer beads. Top to bottom: SRS image (2950 cm^-1^) before sorting; active pixels; SRS image after sorting. **D**. Real-time imaging sorting of live *S. cerevisiae*. Top to bottom: SRS image (2940 cm^-1^) of live *S. cerevisiae* before sorting; active pixels; SRS image after sorting; culturing result of sorted *S. cerevisiae*. **E**. Real-time imaging-sorting of rat brain tissue (thickness 5 µm). Top to bottom: SRS image (2940 cm^-1^) of rat brain tissue before sorting; active pixel; SRS image after sorting. **F**. The same rat brain tissue as E. Top to bottom: SRS image (2850 cm^-1^) of rat brain tissue before sorting; active pixel; SRS image after sorting. **G**. SRS spectra of ejected and unejected spots marked with dashed lines in E bottom.

We further tested the applicability of S-RACE to tissue sections. We prepared a cryosectioned rat brain tissue and ablated multiple regions of interest (ROIs) based on SRS images at 2850 cm^-1^ (**Figure 6E**) and 2940 cm^-1^ (**Figure 6F**). Raman bands centered around 2850 and 2940 cm^-1^ are representative of cellular lipids and proteins and provide contrast in molecular signatures and morphological features of the cryosectioned tissues. Successful tissue ablation was confirmed by SRS spectra (**Figure 6G**), where the ejected spots showed relatively low intensities, while the unimpacted area showed a typical protein-rich Raman spectrum. In the future, instead of single-color SRS, lipid and protein contrast could be combined for tissue ablation with the aid of two comparator circuits. In addition to rat brain tissue, we also tested a bone cancer tissue slice, where 3 FOVs were imaged, featuring protein-rich, vessel, and collagen-rich regions (**Figure S9**). Ejection targets were manually selected, and successful ejections were confirmed by the SRS spectrum after ejection. The size of the ejection spots was quantified. For protein-rich bone cancer tissue, the spot size was 10.98 ± 2.24 μm. The collagen-rich region of bone cancer tissue had a smaller ejection size probably because of the higher rigidity of collagen compared to the protein-rich region. These results highlight the potential of S-RACE as an integrative tool for spatial multiomics measurements across diverse samples ranging from single cells to tissue sections.

## Discussion

Towards the goal of high-throughput label-free single-cell sorting, we reported in this work a stimulated Raman-activated cell sorting system, termed stimulated Raman-activated cell ejection (S-RACE). This platform integrates a multispectral SRS microscope, a laser ejection module, and an online image processing framework. Successful sorting of various samples was demonstrated, including a 1.0 µm polymer beads mixture, bacteria, fungi, and tissue sections. These results illustrate high-throughput sorting of single particles or cells based on their Raman signatures, with precision and minimal impact on the microenvironment around the targeted particle or cell.

S-RACE achieves a throughput of approximately 14 eps on a mixture of polymer beads with 4-color SRS imaging and single-pulse ejection. For 5-color SRS-activated cell sorting, the throughput is approximately 12 eps. Successful qPCR amplification was demonstrated with ejection numbers ranging from 1000-1700, where the ejection can be completed in less than 3 minutes. Considering the sample preparation time, the entire process can be completed in less than 10 minutes. The potential for multiplexed high-throughput sorting is also on the horizon, as different species or cells can be sorted into different wells in the collector for subsequent phenotypic and/or genotypic analyses. The sorting purity has been enhanced by addressing the domino effect of ejection, which can be mitigated by calculating inter-object distances and excluding clustering objects. High sorting yield was achieved for microparticles, microorganisms, and tissue sections, thanks to the high success rate of ejection and the automatic target detection.

Importantly, S-RACE possesses the capability for live cell imaging and sorting, a feat that has long been pursued in Raman-activated cell ejection but remains challenging due to mechanical stress, thermal, and dehydration damage (55). In our study, a thin layer of agarose gel was added to the TiO_2_-coated coverslip to reduce cell damage from the pushing force and heating. This thin and uniform agar layer is crucial to maximize the success rate of cell ejection while maintaining cell viability. A customized receiver with an agarose gel layer was used to protect cells during the landing process. Successful cultivation after ejection for three microorganism species (*S. cerevisiae, C. albicans, E. coli*) was shown. For *E. coli*, a similar recovery rate as the previous report (∼22%) (18) was achieved.

We note that there is still room for improvement in the S-RACE platform. First, LIFT-based cell sorting typically necessitates an air layer to separate the original sample on the top from the sorted cells in the bottom collector. However, this practice contradicts the prerequisites of SRS imaging and consequently attenuates the SRS signal quality. Besides, non-Raman background, e.g. cross-phase modulation, engenders some spurious effects. Removal of the non-resonant background in the SRS image will significantly improve the sensitivity and signal fidelity. The exploration of alternative modalities like frequency-modulation SRS (56), stimulated Raman photothermal microscopy (57) and mid-infrared photothermal microscopy (58, 59) holds promise owing to their high detection sensitivity. The challenges associated with the refractive index mismatch between the sample and the air layer can also be mitigated through the use of photothermal detection regimes. Second, the image dwell time in the current real-time imaging-sorting regime is limited by the relatively low repetition rate of the ejection laser. A higher throughput could be achieved by incorporating a nanosecond laser with a higher laser repetition rate. Furthermore, the use of a hardware-based spectral processor in the module could enhance specificity through multiplex imaging.

We envision that S-RACE would benefit multiple biological applications that were previously challenging or impractical with conventional fluorescence-based or flow-based sorting technologies. First, a synergistic integration of S-RACE with advanced genome, epigenome, and transcriptome sequencing technologies will provide insights into the link between phenotype and genotype at the single-cell level. Second, unlike microfluidics, S-RACE is applicable to tissue samples. Harnessing its high-throughput capacity and advanced image recognition capabilities, discrimination and further genetic analysis of distinct ROIs based on both Raman signatures and morphological features can be achieved. Third, the majority of microbial organisms are still regarded as “dark matter” and waiting to be revealed (60). In tandem with genomic analysis, S-RACE can be a potent tool for discovering unknown species without the need for microorganism cultivation. For example, complex samples from environmental soil could be directly placed onto a TiO_2_-coated coverslip and sorted based on specific phenotypes for single-cell genomics analysis. Tissue samples like sectioned gut tissues could also be studied by S-RACE to gain insights into microbiota activities and the interactions between gut and microbiota while retaining the spatial architecture of the microenvironment. Lastly, it can enable the selection of cells for in vivo cell therapies, including stem cells and chimeric antigen receptor T (CAR-T) cells for personalized medicine. In summary, by integrating an SRS microscope, online image processing, and a cell ejection module, we have developed an automated, high-throughput S-RACE platform for precise single-cell sorting. This platform is compatible with a wide range of samples, spanning from small microorganisms to tissue sections. We expect that S-RACE will open avenues across many biological and biomedical applications.

## Materials and Methods

### Polymer bead mixture

Polystyrene (PS) microbeads with red fluorescence and poly (methyl methacrylate) (PMMA) microbeads with 1.0 μm diameter were mixed with a 2:1 ratio in deionized water. The mixture was dropped onto a coverslip with TiO_2_ coating and then air-dried prior to the S-RACE experiments.

### *E. coli* sample

The *E. coli* strain with GFP label was from the Mary Dunlop lab at Boston University, which harbored a plasmid containing a constitutive promoter-driven superfolder GFP. The cells were first recovered from - 80°C on a Trypticase Soy Agar (TSA) plate for 37°C overnight. Then the TSA plate was stored at 4°C for future use. On the experiment day, a single colony was scrapped from the TSA plate and suspended in a culture tube with 2 ml TSB medium. The suspended cells were cultured at 37°C with shaking at 200 RPM for ∼4 hours.

### *S. cerevisiae* and *C. albicans* sample

The *S. cerevisiae* strain was from the Ahmad (Mo) Khalil lab at Boston University. For both *S. cerevisiae* and *C. albicans* cells, the cells were first recovered from -80°C on a Yeast Peptone Dextrose (YPD) plate for 30°C overnight. Then the YPD plate was stored at 4°C for future use. One day before the experiment, a single colony was scrapped from the YPD plate and suspended in a culture tube with 2 ml YPD medium. The suspended cells were cultured at 30°C with shaking at 200 RPM for ∼4-6 hours.

### *R. glutinis* sample

The *R. glutinis* strain was from the Agricultural Research Service Culture Collection (NRRL). The cells were first recovered from -80°C on a medium No. 6 plate for 30°C overnight. Then the YPD plate was stored at 4°C for future use. A two-phase growing protocol was adapted to promote lipid production. For phase one: a single colony was scrapped from the YPD plate and suspended in a culture tube with 2 ml YPD medium. The cells were cultured at 30°C with agitation at 200 RPM for 24 to 48 hours. For phase two: 100-200 μl of cell culture from phase one was mixed with 2 ml of medium No. 6 supplemented with 3% glucose. The cells were cultured at 30°C with shaking at 200 RPM for ∼96 hours. Medium No. 6 is composed of dextrose 10 g/L, yeast extract 3 g/L, peptone 5 g/L, malt extract 3 g/L.

### Tissue sections

The SJSA-1 tumor tissue and rat brain tissue were fixed with formaldehyde fixative overnight. The tissue section was then transferred to a container with 30% sucrose in 1X PBS at 4°C. After the tissue sank, it was removed from the liquid and embedded in the OCT compound. The tissue sample was then placed in a -80°C freezer until fully frozen. The frozen tissue sample was then sectioned to 5 μm with a cryostat machine (cm1950, Leica).

### S-RACE setup

SRS images were acquired using a lab-built SRS microscope (**Figure S1**). Briefly, a femtosecond laser source (InSight DeepSee, Spectra-Physics) was used for SRS excitation. The laser output two femtosecond pulse trains used for the pump (tunable wavelength) and Stokes (fixed wavelength at 1045 nm), respectively. Both the pump and Stokes beams were chirped for hyperspectral imaging through spectral focusing with high-dispersion glass (SF57, 90 cm in length for the Stokes beam and 75 cm in length for the pump beam). An acousto-optic modulator (522c, Isomet) was used to modulate the Stokes beam at ∼2.5 MHz. A translation stage (Zaber Technologies) was used to scan the interpulse delay between the pump and the Stokes beams, thus the excitation frequency. The combined pump and Stokes beam were directed to a microscope frame by a 2D galvo mirror (GVS002, Thorlabs). The microscope was equipped with a 60X water immersion objective (NA = 1.2, UPlan-Apo/IR, Olympus) or a 40X water immersion objective (NA = 0.8, LUMPLFLN, Olympus). The SRS signal was then captured by a photodiode with a custom-built resonant circuit and extracted by a lock-in amplifier (UHFLI, Zurich Instrument). For the polymer beads sample, the power on the sample was ∼14 mW for 800 nm and 25-40 mW for 1040 nm. For *E. coli*/*S. cerevisiae*/*C. albicans* sample, the power on the sample was ∼14 mW for 800 nm and 50 mW for 1040 nm. For *R. glutinis*, the power on the sample was ∼14 mW for 802 nm and 50 mW for 1040 nm.

For microparticle/cell ejection, a 532-nm laser (ALPHALAS, pulse width 0.89 ns) was collinearly combined with the pump and Stokes before the galvo mirror. The 532-nm laser was operated at a 1.7 A current with a repetitive rate of ∼4 kHz. For automated cell sorting, an acousto-optic modulator (522C-2, Isomet) was used as a pulse picker. A function generator (DG1022Z, Rigol) was used to trigger AOM (∼3kHz, burst mode). The AOM modulation frequency matched the repetition rate of the 532-nm laser. The 532-nm laser was combined with the pump and Stokes beams before the 2D galvo mirror with a 650-nm long pass filter.

### Automatic imaging-sorting

A customized program was developed in LabVIEW that seamlessly integrates SRS imaging, target detection, and single-pulse ejection functions. For single-color SRS images, objects are detected by first generating the object mask and then calculating the centroid of each object. For multi-color SRS images, an additional lease square fitting step was executed, allowing for object classification based on their spectral features. For each targeted object, 2D galvo mirrors were employed to precisely position the green laser on the object. Subsequently, an acousto-optic modulator (AOM) was activated to project the green laser onto the coating. The targeted object was then pushed away from the ejection coverslip and collected by the collector. By moving the sample stage, multiple FOVs were stitched together to achieve high throughput.

Before each day’s experiment, one registration step was performed. This step aligns the coordinates of the SRS image with the position of the green laser.

### The ejection module

Coverslips with TiO_2_ (Titanium dioxide) coating were used for cell ejection. The TiO_2_ coating absorbs the 532-nm laser pulse and forms an ejecting force by nanosecond laser irradiation. Coverslips with TiO_2_ coating were prepared by magnetron sputtering (3.00-inch diameter Angstorm Science ONYX-3 Mag II cathode) with a TiO_2_ target (purity 99.99%, QA13-11200, Angstorm Engineering Inc.). The sputtering time for coverslips used in this work is 2 minutes if not otherwise specified. For live *E. coli* and *S. cerevisiae* sorting, a thin agarose layer was added on top of TiO_2_ by dropping 5 μl 1% agarose and squeezing with another coverslip. This agarose layer was ∼6 μm (measured under a microscope).

For the receiver, a normal coverslip with PDMS (polydimethylsiloxane, thickness ∼200 μm) spacer was used for the polymer beads sample. For imaging GFP labeled *E. coli* in the receiver, a standard coverslip with an agarose gel layer (1%) was used as a receiver and was also with a PDMS spacer (thickness ∼200 μm). For recultivating GFP labeled *E. coli* and *S. cerevisiae*, a petri dish with an agarose gel layer (2% agarose in YPD medium for *S. cerevisiae*, 2% agarose in TSB for *E. coli*).

### Quantitative PCR

For sorted *R. glutinis*, DNA was first extracted using a protocol adapted from a previous study (52). In the final step, DNA in each collection well was diluted to 50 μl. Each qPCR well contained 7.5 μl DNA template, 10 μl PowerTrack SYBR Green Mix, 1 μl forward primer and 1 μl reverse primer (final concentration 1 μM), and 0.5 μl yellow buffer. A 96-well plate with all reagents was then sent to a qPCR machine (StepOne Plus RT-PCR, Applied Biosystems). The qPCR run started at 95°C for 2 minutes, then ran for 40 cycles at 95°C for 15 seconds, 60°C for 60 seconds, and 72°C for 30 seconds. After the qPCR run, a melt curve was measured at 95°C for 15 seconds, 60°C for 60 seconds, and 95°C for 15 seconds.

The primer used in this study was from a previous study (53). The forward ITS2 primer was 5’-GCATCGATGAAGAACGCAGC-3’. The reverse ITS2 primer was 5’-TCCTCCGCTTATTGATATGC-3’. The primers were from Integrated DNA Technologies, Inc.

### Real-time precision opto-control

The comparator circuit box is from The Chi Zhang lab at Purdue University. The comparator box was operated in manual selection mode. The selection threshold was set by turning the knob on the box. The TTL signal output was sent to AOM installed in the optical path of the 532 nm laser. For the TTL signal <0.8 V, the AOM was off. For the TTL signal >2.7V, the AOM was ON and coupled a 532 nm laser into the optical path. Before the experiment, the 532 nm laser was first aligned with the SRS lasers for high-precision sorting.

## Acknowledgments

This work is supported by DOE Grant BER DE-SC0019387 and R35GM136223 to JXC. The SJSA-1 tumor cell was purchased from ATCC and the tissue sample was prepared by Dr. Hongjian He. The rat brain sample was prepared by Mingsheng Li. We appreciate insightful inputs from Professor Michael Wagner at Vienna University and Professor Mary Dunlop at Boston University. We thank Dr. Le Wang, Chinmayee Prabhu Dessai, Zhongyue Guo in the Cheng group, and Dr. Nathan Tauge in Professor Dunlop lab for helpful discussion. We thank Boston University Bio-Interface Technology core facility for their assistance with qPCR experiment and cryosectioning, Boston University Micro and Nano Image core facility for their assistance with confocal imaging, and Boston University Photonics Center Optoelectronic Processing facility for their assistance with magnetron sputtering.

