## Supplemental file for "High-throughput single-cell sorting by stimulated Raman-activated cell ejection"

**This PDF file includes:**

Figures S1 to S9

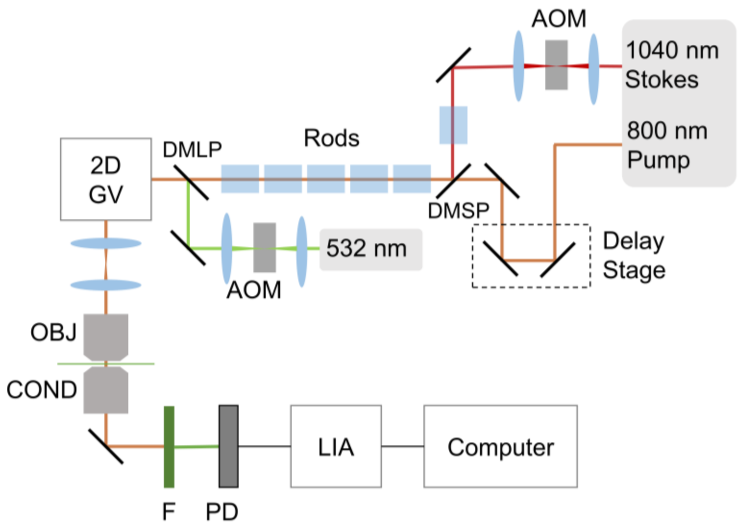


**Fig. S1.** Optical setup for S-RACE. AOM: acousto-optic modulator. DMSP: short pass dichroic mirror. DMLP: long pass dichroic mirror. 2D GM: 2D galvo mirror. L: lens. OBJ: objective. COND: condenser. F: filter. PD: Photodiode. LIA: lock-in amplifier.

A

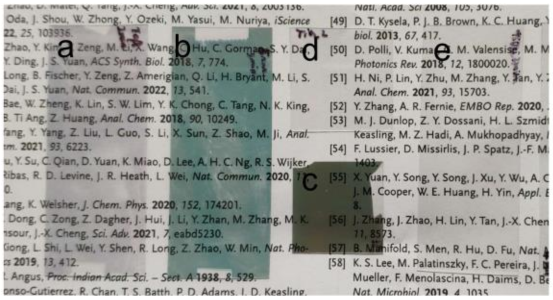


B

| No. | Au (nm) | $TiO_2$ (min) | Ti | SRS | Ejection |
|---|---|---|---|---|---|
| a | 1 | 2 | 0 | Strong background | Good |
| b | 5 | 3 | 0 | Strong background | Good |
| c | 10 | 0 | Unknown | Strong background | Damage |
| d | 0 | 2 | 0 | Good | Good |
| e | 0 | 3 | 0 | Good | Good |

**Fig. S2.** Optimization of laser absorbing coating in terms of SRS imaging and ejection.

**A.** Photo of coverslips with different types of coating.

**B.** Different types of coating. a,b,d,e: fabricated at BU Optoelectronic Processing Facility, parameters as denoted in the table; c, purchased from sigma (serial number 643254-12EA).

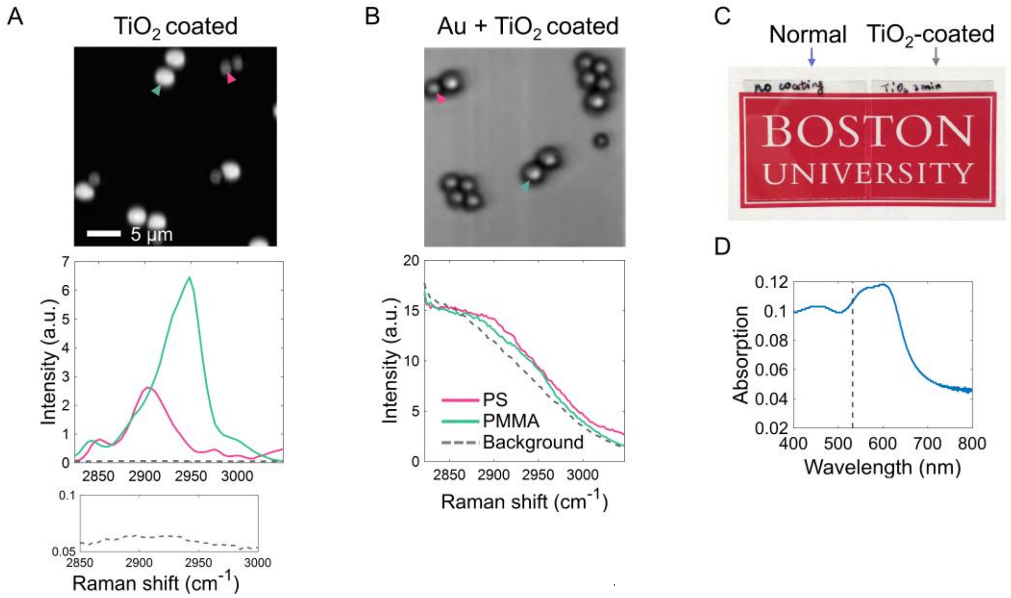


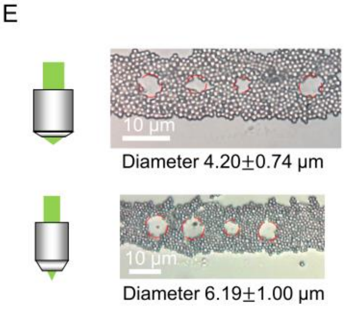


F

| Test | + 1 μm | In focus | - 1 μm | - 2 μm | - 3 μm |
|---|---|---|---|---|---|
| 60X objective | 100% | 93.10% | 81.4% | 36.8% | NA |
| 40X objective; underfilled back aperture | 98.90% | 95.20% | 94.50% | 86.00% | 69.20% |

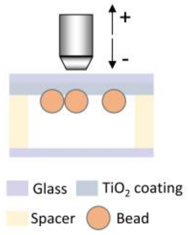


**Fig. S3.** Characterization of the LIFT system used in this work.

**A.** TiO2 coating (2 min sputtering time) has minimal background in SRS image. Top: SRS image of polymer beads mixture (PS 2 μm, PMMA 3 μm); Bottom: SRS spectra of beads marked in the top image (bottom inset shows the background level).

**B.** Au coating (Au 1 nm, TiO2 2 min sputtering time) leads to a high background in SRS image. Top: SRS image of polymer beads mixture (PS 2 μm, PMMA 3 μm); Bottom: SRS spectra of beads marked in the left image.

**C.** Photo of standard coverslip without coating (left) and TiO2 coated coverslip (right) placed on top of a Boston University logo.

**D.** Absorption of TiO2 coated coverslip (2 min sputtering time). The black dotted line denotes 532 nm.

**E.** Demonstration of spatial resolution by microbead cluster (1 μm polymer beads dried on TiO2 coated coverslip). The red circle shows the equivalent diameter of the ejection spots. Left: configuration 1 (60X objective). Right: configuration 2 (40X objective, underfilled back aperture of objective).

**F.** Impact of laser focal position on microbead ejection (1 μm polymer beads).

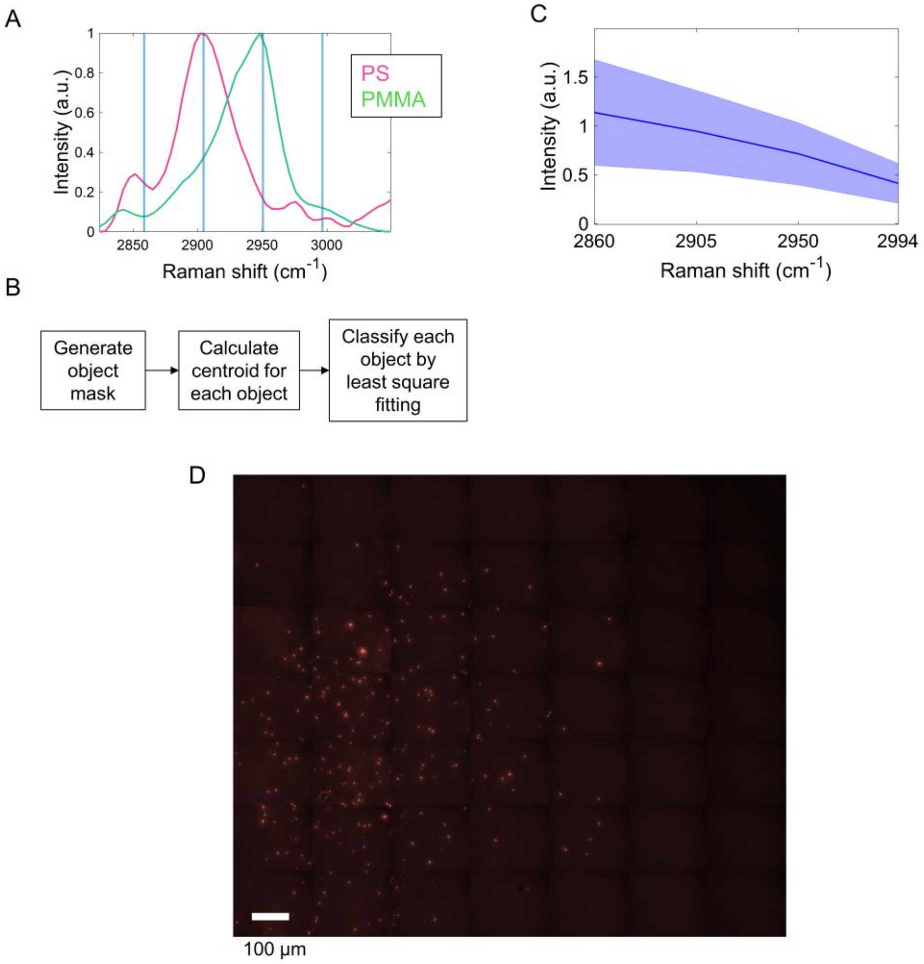


**Fig. S4.** Supporting figure for Figure 2.

**A.** SRS spectra of PS and PMMA beads. Light blue lines represent selected wavenumbers used in Figure 2 (2860, 2905, 2950, and 2994 $cm^{-1}$).

**B.** Steps of single-bead detection used in Figure 2B.

**C.** Multicolor SRS spectra of burning spots in Fig. 2C.

**D.** Wide-field fluorescence image of the collector's bottom. Red dots represent the collected red fluorescence-labeled PS beads after sorting.

| | | PS | PMMA | Yield | Purity | Throughput |
|---|---|---|---|---|---|---|
| exclude PS beads if they are close to PMMA beads (d<2.5 um) | Before | 563 (targeted)<br>108 (clustered) | 273 | 95.2% | 98.0% | 13.9 eps |
| | After | 135 | 262 | | | |
| exclude PS beads if they are close to PMMA beads (d<1.5 um) | Before | 220 (targeted)<br>17 (clustered) | 72 | 88.6% | 93.8% | 11.0 eps |
| | After | 42 | 59 | | | |
| Do NOT exclude neighboring PS beads | Before | 368 | 388 | 90.5% | 76.9% | 6.7 eps |
| | After | 35 | 288 | | | |

**Fig. S5.** Evaluation of S-RACE performance by polymer beads mixture ejection using different bead exclusion conditions.

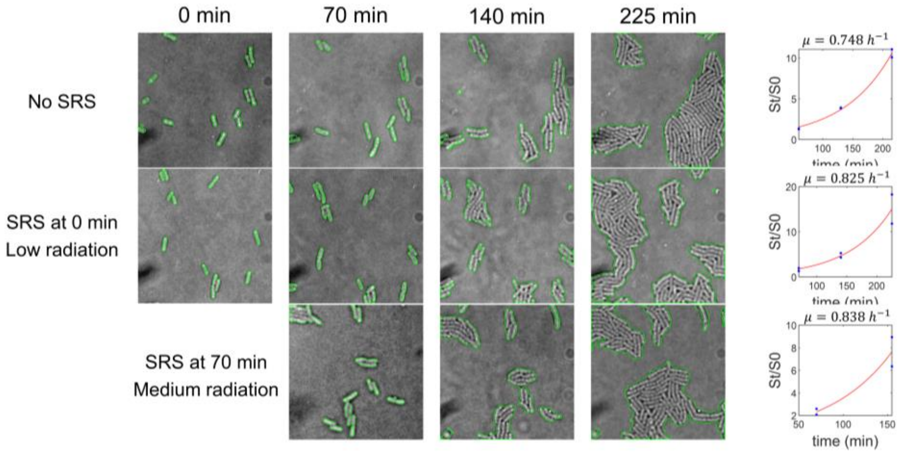


**Fig. S6.** Safe SRS imaging condition has minimal impact on *E. coli* colony growth. Figures in each row represent the time-lapse of *E. coli* colony growing under a specific SRS imaging condition. Top row: no SRS imaging. Middle row: SRS imaging at 0 min. Power: pump beam 24 mW, Stokes beam 50 mW; stepsize: 0.15 µm/pixel; 10 frames. Bottom row: SRS imaging at 70 min. Power: pump beam 24 mW, Stokes beam 100 mW; stepsize: 0.15 µm/pixel; 10 frames. Column in the right, fitting of the cell growth curve for each SRS imaging condition. $\boldsymbol{S}$: colony area (µm$^2$), $\boldsymbol{\mu}$: growth rate (h$^{-1}$), $\boldsymbol{\lambda}$: lag time (h). The equation used in fitting: $\mu = \ln \frac{\frac{S_t}{S_0}}{t-\lambda}$.

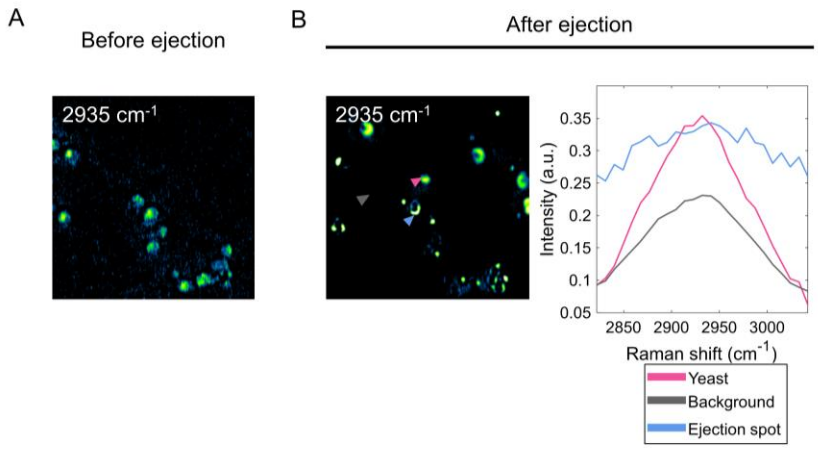


**Fig. S7.** Validation of successful ejection of live cells on an agarose layer.

**A.** Single-color SRS of live *S. cerevisiae* before ejection.

**B.** Left: spectral summation of hyperspectral SRS of the same FOV as A. Right: SRS spectra of ROI denoted by the marks in the left.

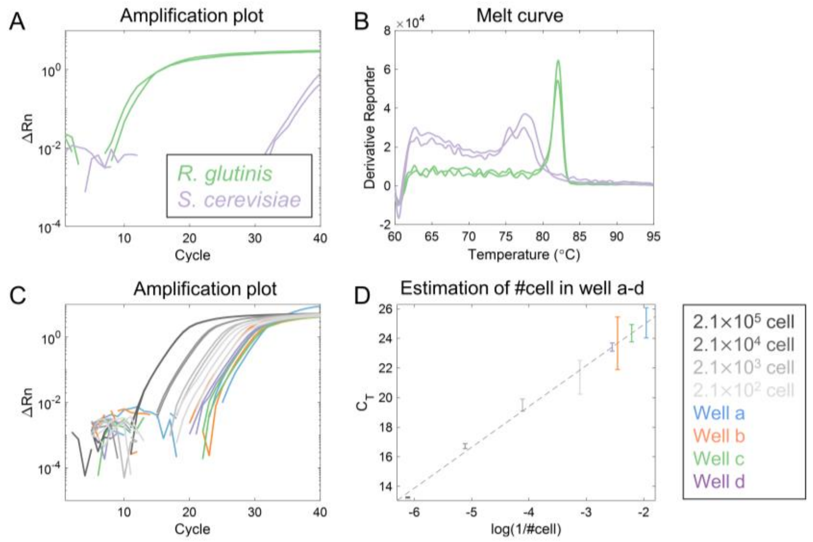


**Fig. S8.** qPCR assay.

**A-B.** Amplification plot and melt curve from pure culture of *R. glutinis* and *S. cerevisiae*.

**C.** Test sensitivity of qPCR assay for detection of ITS2 region in *R. glutinis*. The gray lines denote the DNA extracted from serial dilution of *R. glutinis* pure culture. The numbers in the legend denote the estimated number of cells in the qPCR reaction well. The qPCR amplification plots of the 4 trials in Figure 5 are also shown using the same color scheme.

**D**. Linear fitting of $C_T$ (cycle threshold) and log(1/#cell) of serial dilution samples. The 4 trials in Figure 5H (well a-d) were also plotted with #cell estimated from the linear fitting. The estimated cell number for well a-d are 91, 286, 162, 354. The number of ejection for each small well ranges from 1100 to 1700. Cell number is lower than the number of ejection due to the cell loss during the transfer process and the presence of larger cells that necessitate multiple ejections.

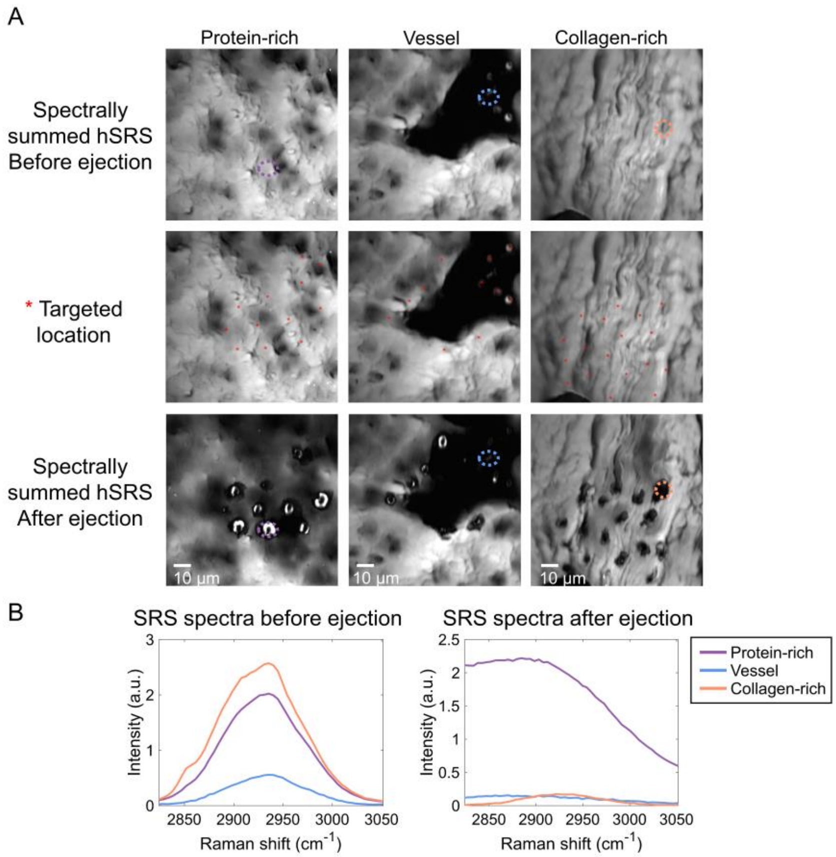


**Fig. S9.** S-RACE of an SJSA-1 tumor tissue section (thickness 5 μm).

**A.** Spectral summation of hyperspectral SRS images of SJSA-1 tissue section before and after ejection. Average ejection spot size: 10.98±2.24 μm (SJSA-1 protein-rich), 7.10±1.35 μm (SJSA-1 collagen-rich). Single-cell ejection was made on single red blood cells.

**B.** Spectra of 3 regions of interest marked by dashed circles in A.
